# Risk-Aware Control for Insulin Delivery via Nonlinear MPC with Safety Barrier Functions and Probabilistic Learning of Uncertainties

**DOI:** 10.1101/2025.09.12.675875

**Authors:** Aliasghar Arab, Seyedreza Mohamadi, Farid Alisafaei, Yashar Mousavi, Katsuo Kurabayashi

**Affiliations:** Department of Mechanical and Aerospace Engineering, New York University, New York, USA; Faculty of Electrical Engineering, Shahrood University of Technology; Department of Mechanical and Industrial Engineering, New Jersey Institute of Technology, 323 Dr. Martin Luther King Jr. Blvd., Newark, NJ, USA; Department of Applied Science, School of Computing, Engineering and Built Environment, Glasgow Caledonian University, Glasgow G4 0BA, UK

**Keywords:** Nonlinear Model Predictive Control, Glucose Regulation, Control Barrier Function, Data-Driven Approximation

## Abstract

Maintaining blood glucose within a physiologically safe range is critical for people with diabetes, as deviations can lead to acute or chronic complications. Hypoglycemia, in particular, represents an immediate threat and requires prioritized mitigation in autonomous insulin delivery systems. This paper introduces a risk-aware hybrid nonlinear model predictive control (NMPC) framework that combines data-driven uncertainty quantification with formal safety assurance through control barrier functions (CBFs). To account for key uncertainties, such as physiological time delays, unannounced meals, and stress-induced glucose variability, Gaussian processes (GPs) are employed as probabilistic estimators. The proposed method dynamically monitors glucose and regulates insulin injection to enforce safe glucose level control by preventing hypoglycemia. The proposed framework is evaluated using validated physiological simulators for various realistic scenarios. The results show a robust performance in maintaining safety under high uncertainty, preparing a foundation for translation into next phase of our research as safe autonomous diabetes management systems.

---

**D** IABETES disrupts metabolism by causing elevated blood glucose levels due to the body’s inability to produce or use insulin effectively. In Type 1 Diabetes Mellitus (T1DM), autoimmune destruction of *β*-cells impairs insulin production, requiring personalized administration to regulate glucose levels and prevent organ damage [1]. Without proper treatment, patients are at risk of hyperglycemia or hypoglycemia, which can cause serious health problems, such as organ damage or coma [2]. Insulin pumps automate glucose regulation by continuously monitoring and adjusting insulin delivery, shown in Fig. 1. Controllers are designed to maintain the glucose level in the accepted range and simplify diabetes management, highlighting the critical need for a safety-guaranteed automated injection system [3].

**Fig. 1.**
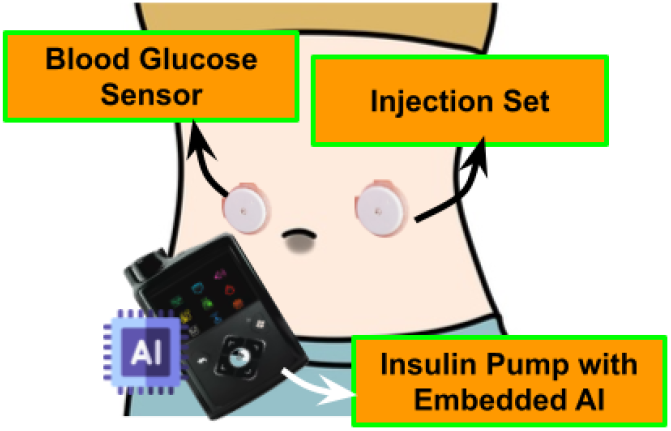
Schematic of an automated insulin injection.

Effective regulation of blood glucose in individuals with T1DM remains a challenging control problem due to insulin delivery delays, meal-related disturbances, and significant intrapatient variability. Previous research has established fundamental performance limitations in glucose regulation stemming from these physiological delays, which inherently restrict achievable control performance [4]. In response to these limitations, efforts have been made to derive performance bounds for optimal insulin infusion, particularly for meals with slow postprandial glucose responses [5]. Moreover, the integration of feedback and feedforward control within model predictive control (MPC) frameworks has shown promise in managing dynamic meal profiles and improving glucose tracking accuracy [6]. More recently, the incorporation of safety layers into MPC architectures, such as interval-based safety constraints and impulsive control strategies, has been explored to better handle patient-to-patient heterogeneity while ensuring safety in artificial pancreas systems [7].

Various methods have been explored for automated blood glucose control and insulin pumps management, such as type 2 fuzzy control [8] and delayed nonlinear model-based strategies [9]. In [10], a higher-order sliding mode control was developed and combined with fuzzy control to handle system nonlinearity. A neural network as a predictive model controller is proposed for glucose control in [11], together with a nonlinear observer-based sliding method to address system uncertainties. In [12], an adaptive observer-based control system was developed using the Bergman minimal model to estimate unmeasurable states. In addition, the Generative Adversarial Network (GAN) architecture is used to improve the effectiveness of the artificial pancreas by generating physical activity and meal information data [13].

More advanced control methods have been developed to address the challenges posed by delayed integration of glucose and insulin into the bloodstream and metabolism. Model Predictive Control (MPC) has been enhanced to better manage blood glucose by predicting insulin-glucose dynamics. In [14], [15], a linear predictive model based on data from 100 patients was used, showing significant improvements in controlling blood glucose levels for individuals with type 1 diabetes. In [16], the combination of feedforward control and MPC further increased prediction accuracy. Furthermore, [17] presented an MPC strategy that used a physiology-based pharmacokinetic / pharmacodynamic model, improving MPC prediction of glucose fluctuations by considering physiological delays and patient-to-patient heterogeneity. This method was validated through in silico simulations and retrospective clinical data. In [18], a dual-hormone MPC system was introduced to administer insulin and glucose in response to exercise with the system equations linearized. Although patient-specific linearized models were designed to enhance the predictive precision of MPC, performance limitations remain due to the inability to fully compensate for the nonlinearity of the models [19].

To address the complexity of glucose regulation, patients were classified by carbohydrate ratios (CR-based models) and grouped by age, with customized linear models for each group. An adaptive Nonlinear MPC (NMPC) in [20] regulated glucose concentration by accounting for insulin’s nonlinear effects but did not explicitly define safe injection ranges. Patient safety was improved in [21] using an ensemble NMPC with a discrete continuous Kalman filter to estimate plasma insulin from glucose measurements. NMPC’s robustness and predictive performance were validated in virtual patients [20], [22], [23], optimizing insulin levels while preventing extreme glucose fluctuations [24]. To address the nonlinear nature of glucose regulation, [25], [26] introduced an insulin infusion advisory system based on NMPC based on NMPC with a personalized metabolic model [27] and data-driven estimators [28]. Evaluated for type 1 diabetes, predictive methods further refined glucose monitoring and control performance and safety.

Advancements, such as MPC frameworks, event-triggered safety layers, and adaptive control methods, have improved glucose tracking accuracy and safety. Despite these advances, previous studies did not fully achieve the prevention of both hypoglycemia and hyperglycemia in type 1 diabetes. More specifically, interval-based safety constraints and impulsive control strategies have been explored to mitigate hypoglycemia risks. But explicit definitions of safe injection ranges and robust safety guarantees are often lacking. It is a critical issue that delayed insulin action and unpredictable disturbances due to meals and stress have prevented existing approaches from precisely regulating hypoglycemia. In terms of hyperglycemia control, dual-hormone MPC systems and physiology-based models have improved glucose regulation. However, they still struggle with intrapatient variability and nonlinearity of glucose-insulin dynamics. This limits their ability to consistently prevent hyperglycemia under varying conditions.

The method proposed in this study fills the existing technical gaps above by integrating CBFs into Nonlinear NMPC and using Gaussian Process (GP) regression models for proactive uncertainty estimation. This ensures robust safety constraints and adaptive control, effectively preventing both hypoglycemia and hyperglycemia.

The remainder of this paper is structured as follows. Section I outlines the problem formulation and the physiological model. Section II-A details the NMPC strategy and integration of the GP model for uncertainty management. Section III presents the experimental evaluation. Finally, Section IV concludes the paper, summarizing key findings and future research directions.

## I. Modeling and Problem Statement

The regulation of blood glucose levels is a critical challenge for individuals with T1DM, requiring continuous insulin control to prevent dangerous fluctuations between hyperglycemia and hypoglycemia. Glucose enters the bloodstream through multiple sources, including meal intake, intravenous infusions, and glucose production by the liver. Although the brain utilizes glucose independently of insulin, other tissues, particularly fat and muscle cells, require insulin to facilitate glucose uptake. Insufficient insulin prevents glucose absorption, leading to hyperglycemia, while excessive insulin can cause severe hypoglycemia, which poses significant health risks. Since insulin degradation directly affects its ability to regulate glucose, an automated insulin injection mechanism must ensure a reliable and adaptive response to changing metabolic conditions. In this section, we formulate the problem as a constrained control problem for a delayed nonlinear system, where glucose concentration should be maintained within safe upper and lower bounds while subject to uncertainties arising from food intake.

### A. Insulin-glucose nonlinear model

To describe the dynamic interaction between insulin and glucose in the human body, we define the nonlinear differential equations representing the balance of glucose *G*(*t*) and insulin *I*(*t*).

#### Definition 1

The term 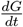 represents the time differential of the net glucose balance, defined as the difference between glucose production and glucose utilization over time.

#### Definition 2

The term 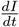 represents the differential equation that describes the net insulin balance, defined as the difference between insulin production and insulin clearance. A delayed nonlinear model for the insulin-glucose system can be presented as a phenomenological version of the Bergman Minimal Model [29] of the glucose-insulin interaction as

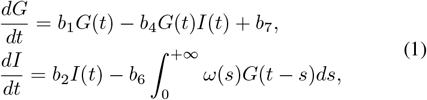

where, the glucose level in the patient’s blood is *G* in *mg/dl* and *I* is the insulin level in the patient’s blood in *µU/ml* with 1 unit (*U*) of insulin is 6.00 nanomoles (nmol). *b*_1_ is insulindependent disappearance rate of glucose, *b*_4_ disappearance rate of glucose-dependent insulin that is concentrated in insulin plasma in (*µU/ml*), *b*_7_ constant amount of glucose increase due to the constant secretion of glucose from the liver, *b*_2_ first order rate of disappearance insulin, *b*_6_ second-order rate of insulin disappearance in (*mg/dl*) per mean plasma glucose concentration per unit time [30]. The kernel *ω*(*s*) is the dynamic selected for the insulin model in the following format.

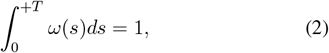

where, *T* denotes the average delay time, which is bounded by *T* < + *∞*. It has been proved in [30] that all solutions obtained from this model converge to a unique, asymptotically stable local equilibrium point, which corresponds to the baseline values of glucose and insulin (*G*_*b*_, *I*_*b*_). Hence, the concentration balance can be considered as follows

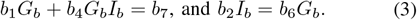

If the delay *T* is small enough, the relative stability of the system is guaranteed. According to the system identification standard based on the intravenous glucose tolerance test (IVGTT), baseline insulin and glucose levels are measured. Then, using the above equation and reducing the number of parameters, identification is performed (i.e., *b*_7_ and *b*_6_ are identified by first determining *b*_1_, *b*_2_, and *b*_4_ along with the measured values *G*_*b*_ and *I*_*b*_). The identification process evaluates the effect of injecting a specific unit of glucose into the veins and its absorption by plasma glucose and insulin in the body over a period of three hours. By setting the time *t* = *t*_0_ as the injection time, the initial conditions can be yielded as

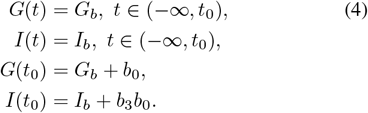

where *b*_0_ is the theoretical increase in glucose concentration after injection and *b*_3_ denotes the initial increase in insulin after injection in (*mg/dl*). *ω*(*s*) can be chosen as [30]

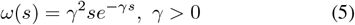

where, *γ* is a time delay constant capturing the physiological delays, including insulin release from *β*-cells and glucagon secretion from *α*-cells in the pancreas. Hence, integrating Eq. (5) with insulin equation in Eq. (1) yields to

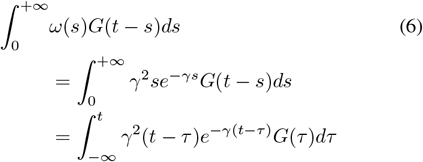

This allows us to write the states of the glucose-insulin model representing delays in the system as

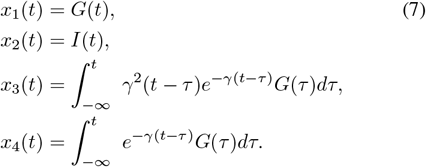

Using linear chain techniques from [31] and the insulin-glucose system in Eq. (1), a 4-dimensional differential equation of the system can be derived as follows

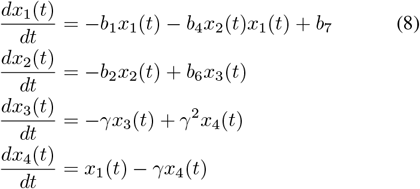

where, using Eq. (4), the initial conditions for the system in Eq. (8) can be written as

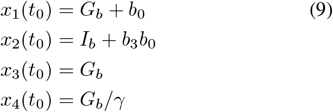

State-space representation of the model can be written as

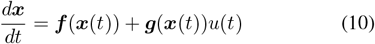

where, ***x***(*t*) is the state vector that includes Eq. (9). *boldsymbolf* captures the dynamics of the glucose / insulin interaction without external input and *boldsymbolg* is part of the system that responds to external insulin input *u*(*t*) injected by an insulin pump. The unified model in (10) represented as

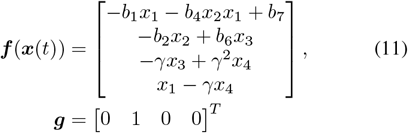

This model represents a comprehensive and unified delayed glucose-insulin nonlinear models, offering a high degree of accuracy in simulating human physiological responses while maintaining a lower complexity compared to previous approaches [31].

### B. Safety constraints for T1DM patients

Equation (11) glucose-insulin model is highly nonlinear and varies between individuals, introducing both nonlinearity and uncertainties into the system. As part of our contribution, we incorporate safety constraints in the form of CBFs into the NMPC framework to address the risks of hyperglycemia and hypoglycemia—offering a new level of safety assurance in glucose regulation.

#### Remark 1

To ensure safety, blood glucose *x*_1_ and insulin injection rate must remain within safe limits. Hypoglycemia must be strictly avoided, while hyperglycemia and injection rate limits are less restrictive.

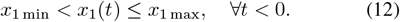

#### Remark 2

The insulin variation rate 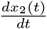 in Eq. (11) should be constrained to clinically safe ranges, typically within 0.5–2.0 units/hour for basal and up to 1–2 units/min for bolus delivery. Exceeding these rates can lead to acute metabolic disruption, hypoglycemia, seizures, or long-term risks such as organ damage and insulin resistance.

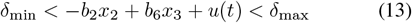

#### Remark 3

Excessive insulin injection must stay bounded as

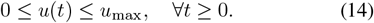

Satisfying these constraints throughout the control design is necessary to guarantee that insulin delivery remains within a safe and effective range.

### C. Uncertainties and Disturbances

The glucose-insulin system in (10) is subject to uncertainties in the model and external disturbances that significantly impact its dynamics. These uncertainties arise from factors such as measurement noise, metabolic variability, and unknown physiological parameters. In addition, external disturbances, including food intake, stress, and exercise, drastically affect glucose concentration, making safe control of insulin delivery challenging.

#### Assumption 1

The glucose-insulin model contains parametric and nonparametric uncertainties, where physiological parameters such as insulin sensitivity, glucose absorption rate, and metabolic clearance vary between individuals and over time. These uncertainties are assumed to be unknown but bounded within a physiologically reasonable range as

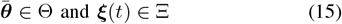

where 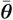 denotes the nominal value of the parameter vector ***θ*** = [*b*_1_, *b*_4_, *b*_7_, *b*_2_, *b*_6_, *γ*], which includes all parameters in (10). These parameters belong to the bounded set Θ, representing the possible values of the parameters constrained within predefined upper and lower limits. In addition, ***ξ***(*t*) represent nonparametric uncertainties, such as unmodeled dynamics, and Ξ characterizes the possible variations in the behavior of the system beyond parametric uncertainties.

#### Definition 3

External disturbances are exogenous inputs that affect the dynamics of the glucose-insulin system but are not directly controlled by the insulin injection mechanism. In this model, disturbances include:

- *Food intake d*_*f*_ (*t*): Introduces a rapid increase in blood glucose concentration as a result of carbohydrate absorption.
- *Stress effects d*_*s*_(*t*): Elevates glucose levels by triggering hormonal responses that increase hepatic glucose production and decrease insulin sensitivity.
- *Exercise: d*_*e*_(*t*) Alters glucose utilization by increasing muscle glucose uptake, which can lower or raise glucose levels depending on intensity and duration.

These disturbances are modeled as uncertain but bounded inputs to the system.

#### Remark 4

Food intake is the most significant external factor that causes glucose fluctuations. Since the amount and composition of food vary daily, an accurate estimation of meal intake is crucial to adjust insulin doses.

#### Remark 5

Stress-induced glucose variations are highly unpredictable due to differences in hormonal responses between individuals. Psychological and physiological stress can lead to increased glucose levels, requiring adaptive control mechanisms to mitigate these effects.

#### Remark 6

Exercise has a bidirectional effect on glucose levels. Although moderate activity typically reduces glucose by increasing insulin-independent glucose uptake, intense physical exertion can cause a temporary increase due to counter-regulatory hormones. This dual effect requires proper estimations and proactive compensations.

A comprehensive model with uncertainties and disturbances is required for a safe control design that must compensate for their effects to ensure glucose remains within a safe range.

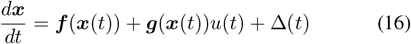

where, Δ(*t*) as the lumped uncertainty and the disturbance represnted in vector format. The lumped uncertainty term Δ(*t*) accounts for both parametric and nonparametric uncertainties, including parametric deviations 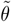, external disturbances *d*_*f*_ (*t*), *d*_*s*_(*t*), *d*_*e*_(*t*), and residual noise *ξ*. It is expressed as

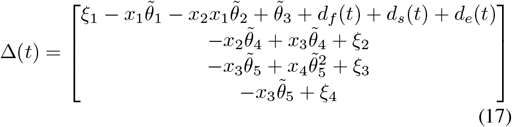

The vector representation of Δ(*t*) is introduced to simplify the design and approximation of future disturbances, which is further detailed in the next section. This formulation enables the proposed predictive control mechanism to proactively compensate for disturbances and ensure that the glucose level remains within safe limits.

## II. Safe Nonlinear Model Predictive Control Design

In this section, a CBF-integrated NMPC controller is designed for safe control of insulin injection to regulate glucose level in T1DM.

### A. Nonlinear Model Predictive Control Framework

This section formulates the NMPC control of insulin injection where CBFs can be integrated as safety constraints. Integration of predictive control with CBF can provide a safer approach to safety-critical control systems, such as glucose-insulin, where safety constraints are directly integrated into NMPC [32]. The architecture of the proposed controller is shown in Fig. 2. The primary objective of the controller is to maintain the glucose level close to a desired level while minimizing oscillations with assured safety. The NMPC optimization problem is formulated as follows.

**Fig. 2.**
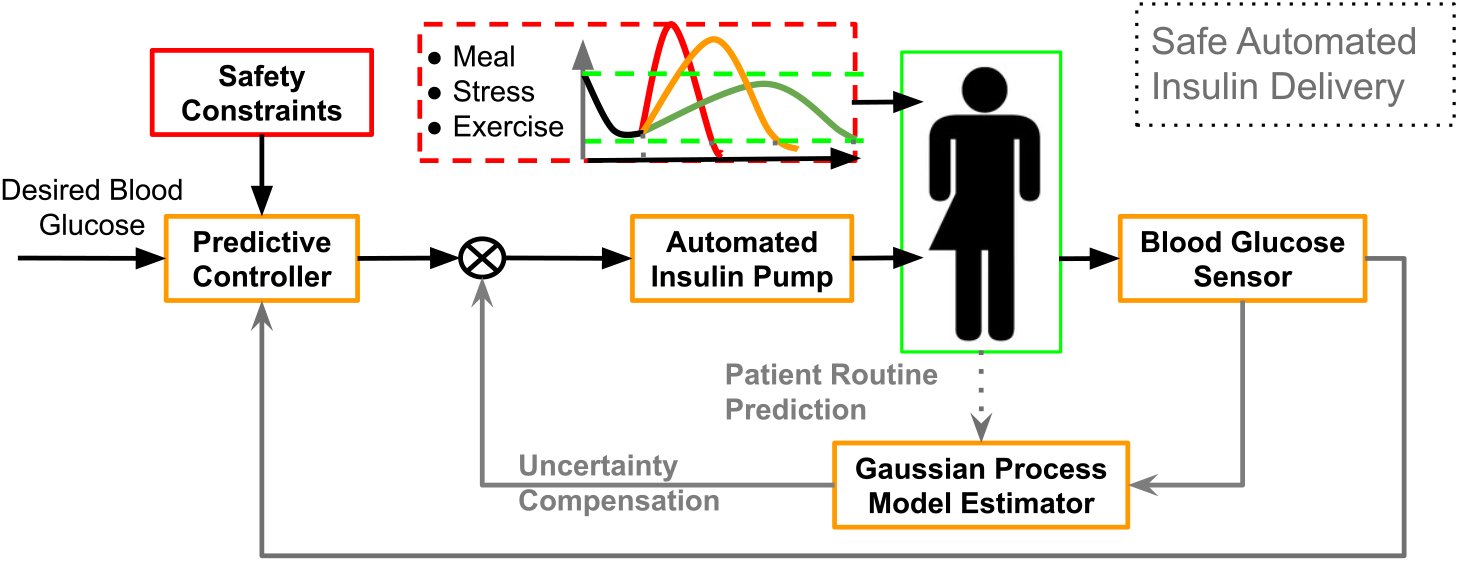
Automated insulin injection with a safe-predictive NMPC, incorporating an adaptive and individualized framework for glucose regulation.

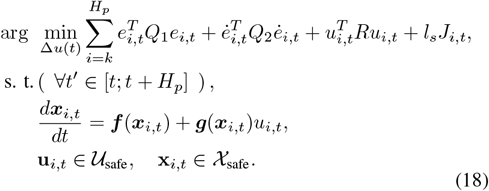

where *H*_*p*_ is the prediction horizon in seconds, *Q*_1_, *Q*_2_, *R*, and *l*_*s*_ are weighting factors and *e*_*i,t*_ is the error value at the *i*th step in the prediction horizon (this notation is used to represent the discrete nature of NMPC for a continuous signal). The error value at a given time *t*, is defined as *e*(*t*) = *G*(*t*) − *G*_ref_ and 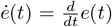 where *ė*_*i,t*_ is the *i*th step of the prediction *ė* within the prediction horizon. The variable *J*_*i,t*_ penalizes the cost function to ensure **x**_*i,t*_ ∈ 𝒳 _safe_ in Remark 1-2 and **u** ∈ 𝒰 _safe_ defined in Remark 3.

### B. Integrating Control Barrier Functions into NMPC

To ensure that the blood glucose level ***x*** remains within the safe set 𝒳_*s*_ for all time *t* > 0, a CBF-based safety restriction as a continuously differentiable function *h*(***x***) is defined that satisfies the following properties

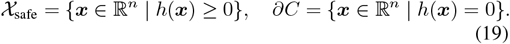

which is positive when *x*_1_ stays within the safe limits. To enforce safety, the control input *u* must satisfy the following CBF condition:

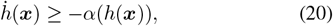

where *α*(·) is an extended class-𝒦 function ensuring forward invariance of the safe set. Using Lie-Brackets and glucose-insulin model with uncertainties in Eq. (16), one can write Eq. (20) as

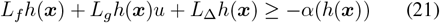

Taking the uncertainties into account for constructing the barrier functions allows us to build a robust safety constraints that can be integrated into the NMPC. Satisfying the Eq. (21) is needed to ensure that the glucose level and the insulin variation rate remain within the prescribed envelope by estimating the upper limits of *L*_Δ_*h*(***x***).

#### Definition 5

Let 𝒳_*s*_ ⊂ ℝ^*n*^ be a safe set. Assume *L*_Δ_*h*(***x***) ≠ 0, ∀***x*** ∈ 𝒳_*s*_. The function *h* is a *Robust-CBF* for NMPC in (18) if there exists *α* ∈ 𝒦_*∞,e*_ such that for all ***x*** ∈ 𝒳 *s*

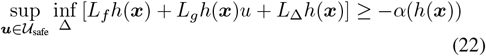

In the wors-case, an uncertain disturbance Δ on the left side of inequality (22) should be considered. We aim to choose a single control input *u* ∈ 𝒰_safe_ for all feasible uncertainties Δ ∈ 𝒟 that satisfy the constraint (21).

### C. Proactive Uncertainty Estimation using GP

GP regression models provide a powerful nonparametric framework for modeling both short-term fluctuations and long-term trends in time series data by leveraging kernels that capture different timescales and periodic behaviors. As a stochastic process where any finite collection of random variables follows a multivariate normal distribution, GPs define a distribution over functions rather than discrete data points. This makes them particularly effective for time-uncertainty modeling in the form of time series, enabling both short-term predictions by capturing transient variations, such as noise and seasonality, and long-term trend estimation by identifying underlying patterns over extended periods. A GP model is trained using glucose-insulin data samples from simulated patient measurements to evaluate the estimation models. The training dataset consists of

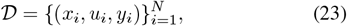

where *x*_*i*_ represents the state (glucose, insulin levels) and the delay models, *u*_*i*_ is the insulin injection rate, and *y*_*i*_ is the measured glucose response. Using a kernel-based learning approach, the GP models Δ(*t*) as:

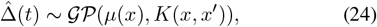

where *µ*(*x*) is the *mean function* capturing baseline dynamics, and *K*(*x, x*^*′*^) is the *covariance kernel function* modeling the correlation between system states. Here, we use the squared exponential kernel, also known as the Gaussian kernel or RBF kernel:

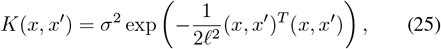

where *ℓ* is the *length scale* parameter controlling the smoothness of the function, and *σ* denotes the *signal variance* controlling the vertical variation. For simplicity, we use an isotropic kernel, meaning that the same length scale *ℓ* is used across all input dimensions. The estimated uncertainty 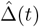 is incorporated into both the safe NMPC controller, where the predicted states allow anticipating where the CBF constraints can reach close to the limit in the prediction horizon for safe control or even notify the patient accordingly.

## III. Implementation and Results

The simulation based validation experiments consists of two distinct phases, 1) focusing on training the GP, 2) evaluating the safety and performance of the system under various test conditions. For this purpose, we define key safety and performance indicators (KSPIs) to evaluate experiments using model-based simulations. Model uncertainties, such as acquiring different types of food, exercising, and stress, are modeled numerically and applied to the simulations to assess the robustness of the controller. The experiment is done in Matlab in both phases.

### A. Learning Phase for Uncertainty Estimations

As mentioned in Section I, the glucose level *x*_1_ is measurable and measured at 5-minute intervals, while the other states of the system cannot be measured directly and should be approximated. Furthermore, the uncertainty value in (16) is not precisely known and the estimation will be taken into account.

In the learning phase, both insulin levels and the patient’s blood glucose concentration are monitored using multi-phase simulations with different sets of disturbances and initial values. NMPC methods allow us to predict *x*_3_ and *x*_4_ using the nonlinear equation (8). To evaluate the proposed method, the blood glucose level of a simulated T1DM patient is controlled by infusing insulin into the body under the regulation of the NMPC controller integrated with CBF safety constraints. This insulin acts as the control signal for the system and is applied as input to the insulin-glucose model. The patient is simulated using an analytical model in which various scenarios such as meal intake (time and amount of carbohydrates), stress events, and initial glucose and insulin conditions are incorporated. These variations are generated using a distribution of uncertainties to emulate realistic and time-varying responses of a T1DM patient. The validation of the method through test simulations is designed to capture a wide range of physiological conditions representative of human patients before testing on a real human. Three times a day, different forms of disturbance in the form of simulated stress are introduced, such as breakfast, lunch, and dinner and three different types, fruits, carbohydrates, and vegetables.

The goal of the controller in the insulin injection system is to keep the patient’s glucose levels at the desired target of 80*mg/dL*, while it should remain in a safe range of 70*mg/dL* to 130*mg/dL*. The disturbance levels are designed to reflect realistic eating patterns, allowing the system to adapt to changes in blood sugar caused by different meal sizes and timings. By simulating glucose fluctuations over a seven-day period, the ability of the system to correct and stabilize these variations is thoroughly tested. In addition, to show the ability of our system in a short time to control the insulin level, the test is carried out in two phases for learning and experiment in 6 hours per day.

### B. Glucose Safe Range as Barrier Functions

The safe glucose level that satisfies the CBF condition in Eq. (20) is defined as

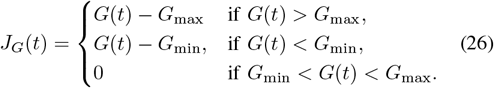

where, *J*_*G*_ is the penalty function for glucose violations as part of the total penalty function *J* (*t*). The control input *u*(*t*), representing the insulin injection rate. The safe range for the glucose level modeled as CBF is incorporated as a hard constraint in the NMPC framework, ensuring that the glucose level always satisfies

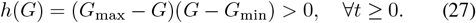

where, *h*(*G*) > 0 satisfies the conditions for a barrier function in Eq. (19). Hence, *G*_min_ < *G* < *G*_max_ and *h*_*G*_ = 0 for *G* = *G*_min_ or *G*_max_. Calculating the time differential of *h*(*G*) in Eq. (27) and putting it in Eq. (21) considering the worst-case scenario for the uncertainties, another safety condition can be written as

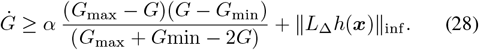

where, this formulation guarantees that insulin injection is optimally controlled while ensuring safety by preventing both hyperglycemia and hypoglycemia. The predictive controller dynamically adjusts insulin delivery in response to disturbances, such as meals, stress, and exercise, while minimizing oscillations in glucose levels.

### C. Implementing Safe NMPC

As described in Section I, the glucose level *x*_1_ is directly measurable every 5 minutes, whereas the other internal states of the system are not directly observable. To effectively regulate insulin delivery under these limitations, the NMPC controller uses estimates of unmeasured states and uncertain parameters. A modified Nonlinear Conjugate Gradient (NCG) method is employed to solve the resulting nonlinear optimization problem [33], [34].

Incorporating CBFs into the NMPC framework requires the control task to be defined as a constrained optimization problem. The cost function aims to keep glucose levels close to the target value, while the constraints, enforced by the CBF, ensure patient safety by limiting insulin injection rates, total insulin dose and maintaining glucose within a safe range. This approach allows the controller to make safe and reliable decisions even in the presence of model uncertainty and partial state observability.

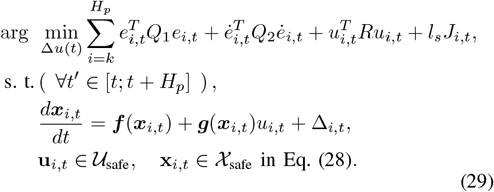

where the safety limits for the CBF penalty function in Eq. (26) are set to *G*_*max*_ = 130 and *G*_*min*_ = 70 as the maximum and minimum limit for the glucose level. When glucose levels are within the safe range, *J*_*G*_(*t*) = 0. However, if glucose levels fall outside the safe range for any given time within the prediction horizon *J*_*i,t*_ > 0, the predictive model will estimate the amount of insulin infusion to ensure glucose stays in the safe range. If the glucose level might exceed the safety limits in the prediction horizon, additional mitigation will be required, such as a notification through smart devices.

### D. Results

Continuous monitoring of glucose levels in automated insulin delivery systems is typically achieved using Continuous Glucose Monitoring (CGM) devices, which often include patch-type sensors. These sensors are attached to the skin, usually in the arm or abdomen, and are minimally invasive, equipped with a small filament inserted under the skin to measure glucose levels in the interstitial fluid. The measured data are transmitted in real time to the insulin injection system or a smartphone unit via a transceiver such as Bluetooth Low Energy (BLE). To realistically emulate this process in our simulation, we discretized the glucose measurement signals and added sensor noise, capturing the sampling characteristics and imperfections inherent to CGM systems.

As shown in Fig. 3, blood glucose levels during the experimental phase were measured and maintained at approximately 80 mg/dL, which was designated as the target value throughout the day. The system consistently regulated glucose around this desired level, even in the presence of external disturbances such as meal intake. Meal-induced perturbations caused transient deviations in glucose levels; however, the NMPC effectively brought glucose back within safe limits, adhering to predefined safety restrictions. The extent of these deviations varied between meals, with the most significant observed at lunchtime due to the highest food intake. NMPC responded by infusing insulin to reduce the glucose level from a peak of approximately 140 mg/dL to the desired 80 mg/dL. Despite substantial disturbances caused by meal consumption, the insulin infusion process successfully stabilized glucose levels.

**Fig. 3.**
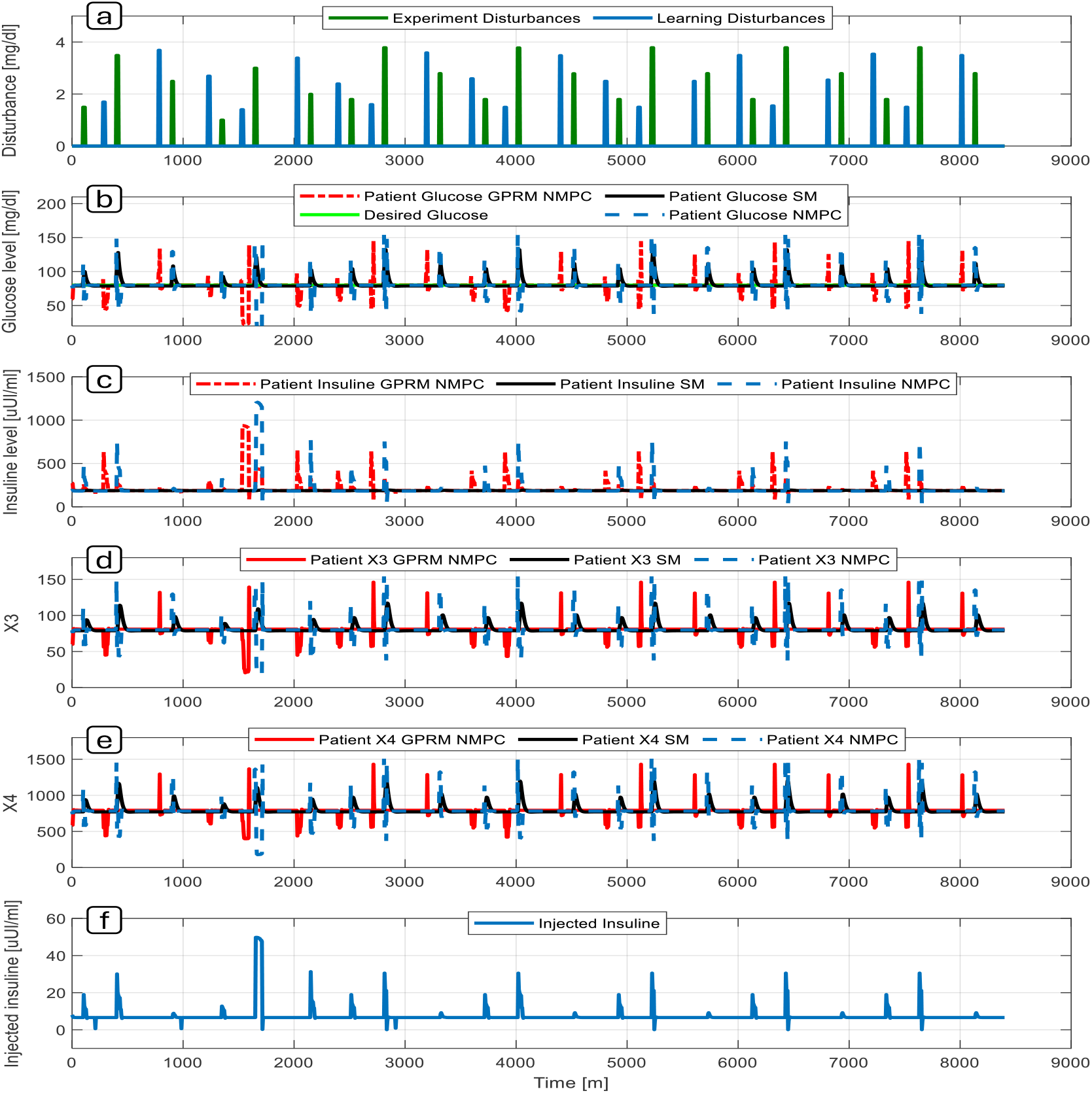
Automated insulin injection in seven days: a) Different disturbances by food intake in seven days during the learning and experiment stages, b) Glucose level control under disturbances, c) Insulin level control using GP NMPC, d) NMPC third state (*x*_3_), e) NMPC fourth state (*x*_4_), f) Injected insulin by NMPC. All data are based on simulations using analytical models implemented in MATLAB.

**Fig. 4.**
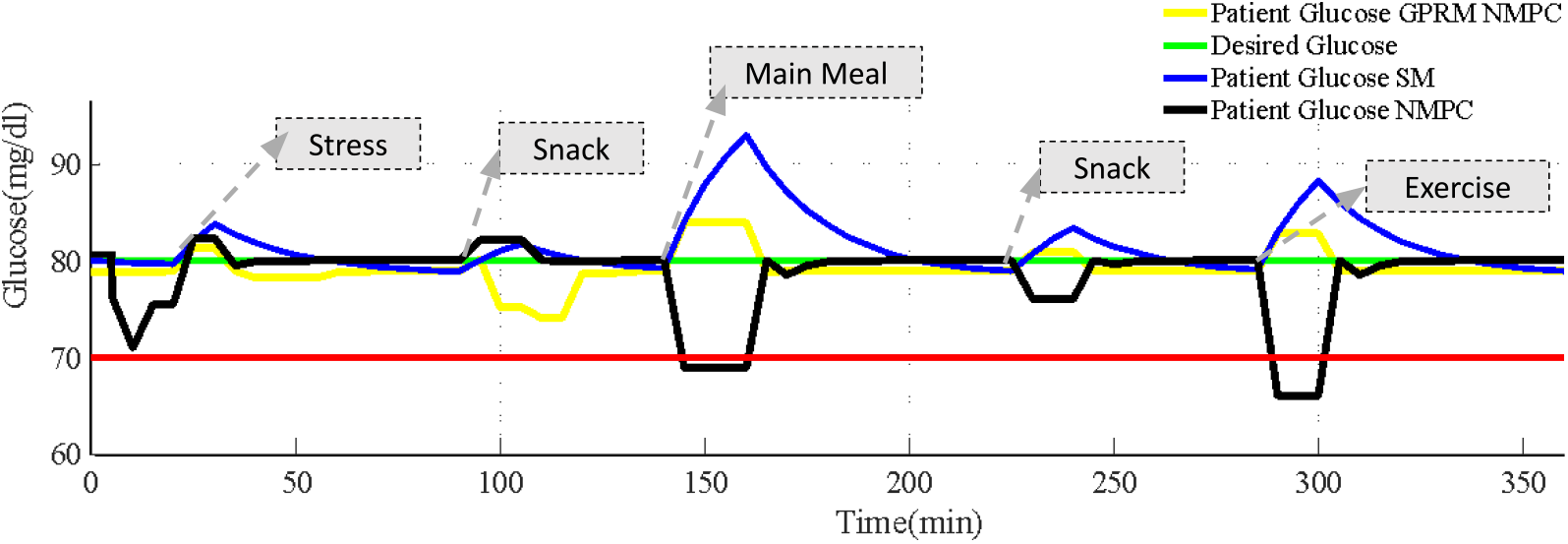
Automated insulin injection in 6 hours (360 Mins) where injected insulin by GPRM NMPC mitigates the risks of surpassing safety limits. The robustness and response speed of the proposed method are also compared to evaluate its performance under variable conditions.

For the remaining meals of the day, breakfast, and dinner, the system exhibited comparable behavior, albeit with less pronounced disturbances than those observed during lunch. This pattern is evident, which demonstrates smaller deviations and a more rapid stabilization by the NMPC. The states of the system, representing the internal physiological dynamics, were also estimated by the GP [35]. Fig. 5 shows the trained GP model with the trained uncertainty range to estimate Eq. 17. In particular, the estimation error for insulin levels was lower than that for glucose, indicating better performance of the GP model for this parameter. The general consistency across all estimated states highlights the robustness of the GP in modeling uncertainties within the safe GPRM NMPC framework compared to other methods such Sliding-Mode (SM) and NMPC without GPRM uncertainty approximation.

**Fig. 5.**
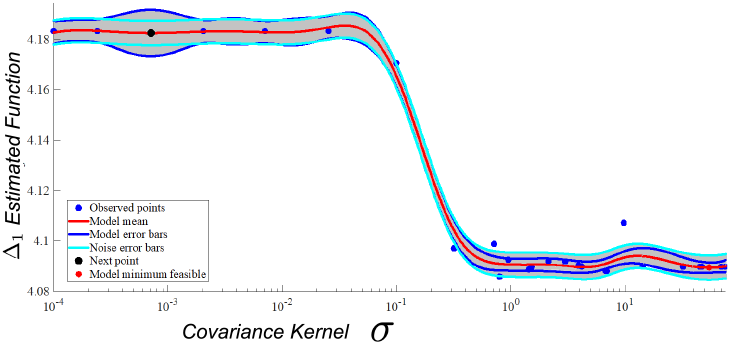
Approximation of Glucose Uncertainty with upper and lower range using GPRM.

#### 1) Robustness and Safety Evaluation of Safe NMPC

In the experimental phase, the learned functions of the GP were integrated into the NMPC controller to compute the cost function and guide the control decisions. This phase was designed to test the controller’s *robustness* by introducing disturbance profiles that differed significantly from those used in the training phase, thus evaluating the controller’s ability to generalize. As shown in Fig. 3-a, the magnitude and timing of disturbances during the experimental phase varied substantially from those in the learning phase. Despite these differences, NMPC, using the GP’s learned models, maintained glucose concentrations close to the desired set point of 80 mg/dL, thus demonstrating the robustness of the system in the face of unexpected fluctuations. The system effectively managed variations in disturbance magnitude, particularly during longer meal periods, with minimal deviations from the target glucose level. The comparison results of the Safe NMPC, standard NMPC, and sliding mode (SM) control methods are illustrated in Fig. 3-b.

A key capability demonstrated during this phase is the system’s operation in the absence of continuous sensor input. In real-world settings, patients may not have access to continuous glucose monitoring due to sensor limitations or cost. Therefore, a model capable of estimating and regulating glucose based on previously learned patterns is critical to ensuring safety. As shown in Fig. 3-b, the Safe NMPC, supported by the GPRM estimator shown in Fig. 5, maintained effective glucose regulation even with variations in meal sizes and timings. Its ability to handle both overfeeding and irregular meal schedules highlights its adaptability and practical utility. In particular, during large glucose spikes, such as those observed at lunchtime, the Safe NMPC maintained glucose levels within predefined safety limits, a critical characteristic to ensure patient health. These findings validate the system’s capacity to compensate for differences between training and deployment scenarios. The *safety* of the system can be quantified using the Safety Performance Index (SPI), defined as

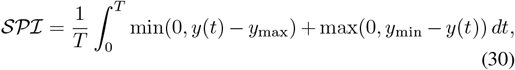

where *y*(*t*) is the glucose concentration at time *t*, and *y*_min_ and *y*_max_ define the safe operating bounds. A more negative SPI value indicates better safety performance.

Similarly, the system’s *robustness* to dynamic disturbances is characterized by the Robustness Index (RI), defined as:

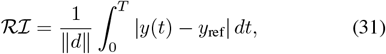

where *d* denotes the disturbance profile and *y*_ref_ is the reference glucose level. Lower values of this integral indicate less deviation from the reference under disturbance, hence higher robustness.

As shown in Fig. 3-c, the insulin usage across the three control strategies is compared. The Safe NMPC approach achieved tighter glucose regulation with reduced insulin injections, outperforming both the NMPC and the SM approaches. This efficiency reduces the risk of hypoglycemia due to insulin overdose, while also improving patient outcomes. Figures 3-d and 3-e show the third and fourth states of the system, representing the internal glucose-insulin dynamics. Figure 3-f shows the insulin infusion profile administered by Safe NMPC during this phase.

Performance metrics are summarized in Table I, which presents the SPI, RI, and convergence time for each controller. The Safe NMPC achieved the lowest SPI of −0.27, outperforming the Iterative Feedback Linearization (IFL) controller [36] with SPI of −0.04, and the CBF controller [37] with SPI of −0.23, demonstrating superior safety. The Safe NMPC also achieved the highest robustness index (RI = 2.89), indicating enhanced resilience to nonlinear disturbances, compared to the CBF controller (RI = 2.44). Furthermore, the convergence time of 3.3 seconds achieved by the Safe NMPC was faster than that of SIFL (3.5 s) and CBF (3.6 s), allowing faster response to disturbances.

**TABLE I.** Safety, robustness and performance comparison.

| Control Method | $SPI$ | $RI$ | Convergence time (sec) |
| --- | --- | --- | --- |
| SIFL | -0.04 | 1.72 | 3.5 |
| Proposed NMPC | -0.27 | 2.89 | 3.3 |
| Basic CBF | -0.23 | 2.44 | 3.6 |

## IV. Conclusions

This work addresses the challenge of safely regulating glucose levels in T1DM under unpredictable disturbances such as meals, stress, and exercise. We proposed a safe NMPC framework that incorporates CBFs and GPs to predict and compensate for individual-specific uncertainties while enforcing safety constraints. Simulation results tested on demonstrate that our controller maintains glucose levels near the desired target of $80 mg/dL$ and outperforms standard NMPC and sliding-mode controllers in the presence of varying disturbances. The integration of CBFs and GPs into NMPC offers a robust and scalable approach for artificial pancreas systems and other automated medical technologies. Future clinical studies are needed to validate the safety and effectiveness.

---

**Aliasghar Arab** (Senior Member, IEEE) Aliasghar Arab received the B.Sc. degree in robotics engineering and the M.Sc. degree in electrical engineering from Shahrood University of Technology, Iran, and the Ph.D. degree in mechanical and aerospace engineering from Rutgers University, NJ, USA, in 2021. He was a Robotics Research Scientist at Nokia Bell Labs and Verizon, a Postdoctoral Researcher at North Carolina A&T State University, and is currently an Adjunct Professor at NYU Tandon School of Engineering and Director of the Agile Safe Autonomous Systems (ASAS) Laboratories. He also leads the R&D team at GenAuto.AI. Dr. Arab has received innovation awards and grants from the National Science Foundation (NSF) and the New Jersey Economic Development Authority (NJEDA). His research interests include safety-critical and nonlinear control, model predictive control, motion planning, generative AI for robotics, and quantum-safe systems. He serves on UL and IEEE standard technical panels and is an active reviewer for IEEE journals and conferences in robotics, artificial intelligence, and control. He is a member of the American Society of Mechanical Engineers (ASME), the Society of Automotive Engineers (SAE), and the Association for the Advancement of Artificial Intelligence (AAAI).

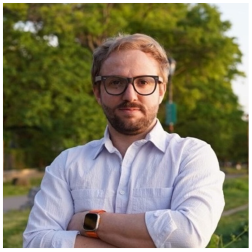

**Sayereza Mohamadi** Sayed Reza Mohammadi received the B.Sc. and M.Sc. degrees in robotics and mechatronics engineering from the Department of Electrical and Robotic Engineering, Shahrood University of Technology, Iran. He was recognized as a top researcher and received the Graduate Excellence Award from the university’s Center of Excellence. His academic background focuses on intelligent robotic systems, mechatronic design, and autonomous control.

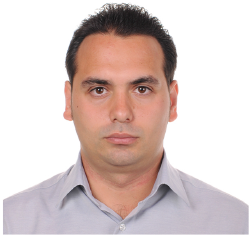

**Yashar Mousavi** (Member, IEEE) Yashar Mousavi received the Ph.D. degree from Glasgow Caledonian University, Glasgow, U.K., in 2023. He is currently a Senior Analytical Engineer with American Axle & Manufacturing, Detroit, MI, USA. He is majored in artificial intelligence-based modeling, simulation, and analysis of combustion-based and electric vehicles’ power transfer units, gear shift actuation systems, eLocker electric drive units, and full-vehicle driveline systems. He has carried out various projects for top-notch companies, such as General Motors, Stellantis, Ford, Mercedes Benz, and Volkswagen. He is also a Controls Integration Researcher with the Power and Renewable Energy Systems (PRES) Team, Glasgow Caledonian University. His research interests include control theory and applications, vehicle dynamics modeling and analysis, artificial intelligence, power systems analysis, robust nonlinear control, fault-tolerant control, renewable energy, and robotic systems modeling and control. He is a member of the IEEE Young Professionals. He served as a reviewer and the guest editor for numerous journals and conferences.

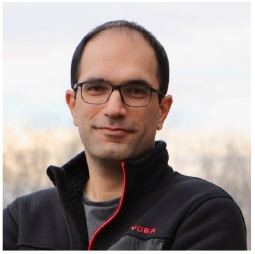

**Farid Alisafaei** Farid Alisafaei is an Assistant Professor of Mechanical and Industrial Engineering at NJIT. His research in mechanobiology focuses on how mechanical forces shape physiological and pathological processes, including wound healing, fibrosis, and the progression of diabetes. As a former NIH-T32 postdoctoral fellow at the University of Pennsylvania and a member of the NSF Center for Engineering Mechanobiology, Dr. Alisafaei has developed integrated computational and experimental approaches to investigate how physical changes in the extracellular environment affect cellular functions, including gene expression and differentiation. He leads the Mechanobiology and BioMechanics (MBBM) Lab, where his research explores the biomechanical pathways that drive disease states. While the present study focuses on the use of control barrier functions and Gaussian processes for safe insulin-glucose regulation, Dr. Alisafaei’s mechanobiological work offers a complementary foundation for understanding how mechanical and systemic factors influence metabolic disorders like diabetes, paving the way for more comprehensive and physiologically grounded control strategies.

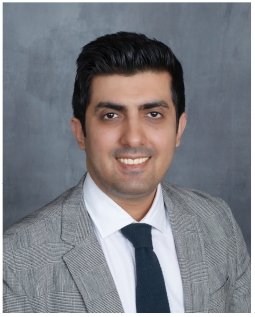

**Katsuo Kurabayashi** (Senior Member, IEEE) Katsuo Kurabayashi received the BS degree in precision engineering from the University of Tokyo, in 1992, and the MS and PhD degrees in materials science and engineering from Stanford University, Stanford, CA, USA, in 1994 and 1998, respectively. He is currently a professor and the chair of mechanical and aerospace engineering with New York University, Brooklyn, NY, USA, and a former Professor of mechanical engineering with the University of Michigan, Ann Arbor, MI, USA.He has authored and co-authored 180 peer-reviewed papers, including Advanced Materials, Nature Communications, ACS Nano, and Nano Letters. He holds 11 U.S. patents. His research interests include optofluidics, nanoplasmonic and biomolecular biosensing, and BioMEMS/microsystems for immunoassay, clinical diagnosis, single-cell study, and analytical chemistry. He was the recipient of 2001 NSF Early Faculty Career Development (CAREER) Award. He is also a Fellow of the Royal Society of Chemistry (RSC) and the American Society of Mechanical Engineering (ASME).

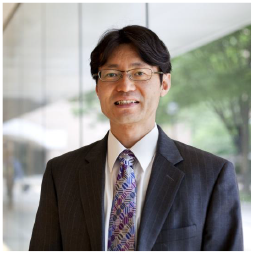

